# A Signaling Network based Computational Model to Uncover Loop as the Novel Molecular Mechanisms for Medulloblastoma

**DOI:** 10.1101/561076

**Authors:** Jielin Xu, Fuhai Li

**Affiliations:** Institute for Informatics (I2), Washington University School of Medicine, Washington University in St. Louis, St. Louis, Missouri, United States; Department of Pediatrics, Washington University School of Medicine, Washington University in St. Louis, St. Louis, Missouri, United States

**Keywords:** loop, drug resistance, molecular mechanisms, signaling network, medulloblastoma

## Abstract

**Motivation:** Medulloblastoma (MB) is the most common malignant brain tumor in children. Despite aggressive therapy, about one-third of patients with MB still die, and survivors suffer severe long-term side effects due to the treatments. The poor post-treatment outcomes are tightly linked to unpredictable drug resistance. Therefore, before developing robust single drug or drug combination recommendation algorithms, uncovering the underlying protein-protein interaction (PPI) network patterns that accurately explain and predict drug resistances for MB subtypes is essential and important.

**Results:** In this study, we hypothesize that the loop sub-structure within the PPI network can explain and predict drug resistance. Both static and dynamic models are built to evaluate this hypothesis for three MB subtypes. Specifically, a static model is created to first validate that many reported therapeutic targets are located topologically on highly deregulated loop sub-structure and then to characterize the loop for tumors without treatment. Next, with the after-treatment time-series genomics data, a dynamic hidden Markov model (HMM) with newly designed initialization scheme estimates the successful and unsuccessful occurrence probabilities for each given PPI and then re-delineates the loop for post-treatment tumors. Finally, the comparison of loop structures pre- and post-treatment distinguishes effective and ineffective treatment options, demonstrating that the loop sub-structure is capable of interpreting the mechanism of drug resistance. In summary, effective treatments show much stronger inhibition of cell cycle and DNA replication proteins when compared to ineffective treatments after considering the cross talk of multiple pathways (the loop).

## Introduction

MB is the most common malignant childhood brain tumor which are mostly found in patients with age under 16. There are four subtypes of MB: WNT, SHH, Group 3 and Group 4 [1]. For children with average-risk and high-risk MB, the 5-year survival rates are 70%-80% and 60%-65% [2], respectively. Therefore, about one-third of children diagnosed with MB will die of the disease. The major reason for the failure of provided treatments is unpredicted drug resistance. Various drug-resistance mechanisms are generally classified as either intrinsic or extrinsic. Extrinsic factors are mostly related to interactions with stromal cells and other tumor cells while intrinsic factors are mainly genetic variations, epigenetic altering and cross talk between multiple pathways. Genetic variations can lead to drug resistance as the first reasonable explanation by reducing the activities of drugs, for example, down-regulation or mutation in proteins on the same pathways as the drug targets [3]. Mutations of the target itself can also reduce or completely alter drug functions and lead to drug resistance [3]. The second type of intrinsic drug resistance mechanism is called multi-drug resistance. Multiple drugs with different mechanisms for entering the cell are used for effective chemotherapy since cell surface receptors or drug transporters can be lacking. However, cancer cells often present various unexplained simultaneous resistances to functionally unrelated drugs, a phenomenon that defines multi-drug resistance. The effects of multi-drug resistance would finally either block the cell apoptosis process or activate the cancer cell DNA repair process which serves as the third intrinsic drug resistance mechanism [4], [5]. The blocked cell death pathway can also result from up-regulation of cell survival promoting proteins, e.g., BCL2, or its upstream PI3K-AKT signaling pathways and down-regulation of cell apoptosis promoting proteins e.g., BAX. [3]. Epigenetic alteration (DNA methylation and histone alternations) is the fourth factor which could cause drug resistance. DNA methylation can inhibit tumor suppressor genes and also amplify the expression of oncogenes via hyper-methylation. Cross-talk between multiple pathways has been reported to induce resistance in various types of tumors, e.g., for breast cancer [5].

In addition, mathematical computational models have been developed to study and simulate cancer drug resistance. The first group of models are based on various types of classical differential equations. Faratian et al. [6] applied an ordinary differential equation (ODEs) based kinetic model to study the behaviors of PI3K in RTK inhibitor resistance. Sun [7] et al. developed a kinetic model to study the cross talk between EGF/EGFR/Ras/MEK/ERK and HGF/HGFR/PI3K/AKT pathways, and their model verified the connection between cross talk of several pathways and drug resistance. Coldman et al. [8] used a stochastic model to simulate the initialization and development of drug resistances due to point mutation and ultimately to estimate the probabilities of drug sensitive and drug resistant cells under different treatment protocols. Sun et al. [7] utilized lists of stochastic differential equations (SDEs) to simulate and differentiate the dynamics of drug-sensitive and drug-resistant cells, and their model presented distinct synergy patterns for BRAF + MEK versus BRAF + PI3K combos. Data-driven models are also widely applied to study drug resistance. Shukla et al. [9] analyzed methylation profiles of glioblastoma (GBM) samples and implemented a cox regression model to identify methylation biomarkers, concluding that targeting the NFkB pathway could prevent drug resistance for GBM. With patients’ expression data, Zeng et al. [10] developed a time involved module network rewiring (MNR) model to prove that functional connections and reconnections for consistent modules could be taken as biomarkers to measure drug efficiency and resistance. In addition to different equations, the time-series HMM [11] model should also be capable of simulating post-treatment tumor dynamics. The expectation maximization (EM) algorithm [12], designed for likelihood maximization with latent variables, serves as the corresponding numerical strategy. However, “effective” initialization is always difficult when encountering the HMM algorithm due to the nonconvexity of the objective likelihood function. Starting with many different random initializations could always approach the global maximum likelihood numerically, however, it is time-consuming and could lead to very unstable dilemma with quite different initialization parameters and quite similar likelihoods - a definitely unacceptable situation, especially for our case. Several clustering related algorithms [13], [14] for continuous observations have proven their capabilities in finally arriving at the global maximum of likelihood. However, for discrete observations, it is usually hard to find relative robust and stable initialization schemes.

Our loop hypothesis has firstly been proposed in [15] for LKB1 loss non-small cell lung cancer, and one PI3K and mTOR dual inhibitor based on the loop hypothesis has been numerically filtered out and biologically validated as the effective treatment. In this work, we will present a more detailed explanation about the hypothesis and its numerical validation for drug resistance. We hypothesize that the loop sub-structure, which considers the cross talk of multiple pathways (all signaling and cellular growth and death pathways) inside the protein network, covers promising therapeutic targets and concurrently could explain and predict drug resistance. The paper is organized as follows: The **Materials and Methods** section explains the details of both static and dynamic models. The **Results** section first starts with a static model and then the outputs from the dynamic model will be checked both mathematically and biologically to guarantee their further engagement in explaining and validating the loop theory. Next, we present the most detailed explanation about why and how to utilize the loop to characterize drug resistance with multiple MB subtypes and treatment protocols. After validating and explaining the loop theory, interesting contrasts and connections between pre- and post-treatment loops are discussed. Discussions and conclusions are consequently summarized.

## Materials and Methods

### Gene expression data and background signaling network

In order to prove our hypothesis, for the static model, we use gene expression data of 194 MBs and 11 normal cerebellums from Cho et al [16]. In Cho’s work, the RNA expression data of all 194 MBs are clustered into 7 subgroups: C1 subgroup represents MYC-driven MBs, C3 subgroup represents SHH type MBs, and C6 subgroup covers WNT type MBs. Three independent time-series RNA expression data are used as the input for the dynamic model. RNA expression data of time course experiments of OTX2 silencing and control experiments in the D425 MB cell line, characterized as representative of Group 3 MBs [17], are provided in Gene Expression Omnibus (GEO) with series GSE22875 with 6 independent time points which form 729 time-sequenced data samples. Time-series RNA expression data for effective SMO inhibitor MK-4101 on SHH type allograft MB tumors can be download from GEO with series number GSE77042 (two doses, 40 and 80 mg/kg each day with five independent time points, forming 720 and 960 time-sequenced data samples, separately). Different categories of pathways for system biology research represented in the Kyoto Encyclopedia of Genes and Genomes (KEGG) pathway database [18] are selected as directed background molecular interaction networks, and we focus particularly on signal transduction and cell growth and death pathways for humans. The R package “Pathview” [19] is applied to download KGMLs for all humans’ pathways while the package “KEGGgraph” [20] facilitates the extraction of node connections from KGMLs for each pathway. Except for activation and inhibition PPIs, the PPIs (phosphorylation, dephosphorylation, ubiquitination, deubiquitination, glycosylation, methylation, indirect effect or state change) are classified as either activation or inhibition according to their specific functional behaviors for provided different protein pairs. The static and dynamic statuses of the remaining two PPIs, binding/association and dissociation, together with activation and inhibition PPIs, are defined in the following sub-section independently.

### Pre-treatment loop extraction model (static model)

For a specific pathway in the KEGG pathway database, a complete full subroutine is a sequence of protein family interactions starting from receptor protein families on the cell membrane down through the terminal protein families in the nucleus or cytoplasm. For each provided subroutine in KEGG, we construct 2 binary sequences from the backward direction with the first sequence always ending with 1 and the second one always ending with 0. Assuming a subroutine with length L, then for the following edges, the binary input is described in the following pseudo-code form:

~~~
Loop each index from L−1 to 1 (decreasing order)
 If current relation is not inhibition/repression:
  binary input [current index] = binary input [current index + 1]
 else:
  binary input [current index] = 1 − binary input [current index + 1]
~~~

It is worth mentioning that the binary sequence is defined with respect to edges, i.e., a subroutine with L + 1 nodes has length L, and, therefore, each subroutine is equipped with 2 binary sequences with both lengths equal to L. For each given PPI, we define 2 edge scores: state_1_score and state_0_score, which describe the probabilities of successful and unsuccessful occurrence of given PPI, separately. For the static mode, the definitions of these 2 scores are characterized as follows. The network extraction model in our numerical experiments is applied to all 3 MB subtypes. For a specific gene A, the expression ratio is defined as:

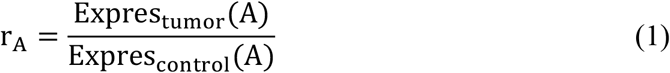

Where *Expres_tumor_*(*A*) and *Expres_control_*(*A*) denotes the average expression of gene A among tumor and control samples. In our edge score model, assuming A and B are two proteins, we always assume protein A sits on the left of the edge and protein B sits on the right of the edge. We use the following equation to characterize the probability of occurrence of type I relation (activation or expression):

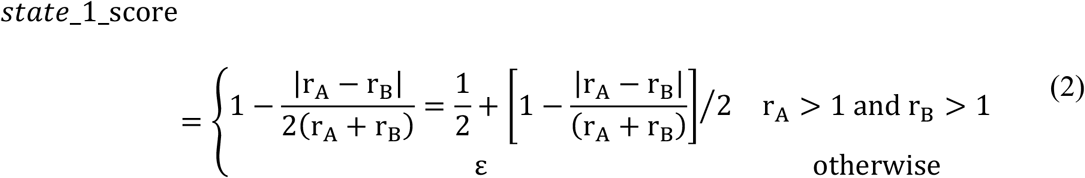

For type II relation (inhibition and repression), we have:

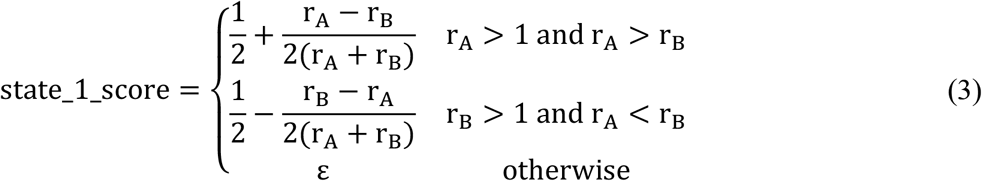

For type III and IV relations (binding/association and dissociation), we have:

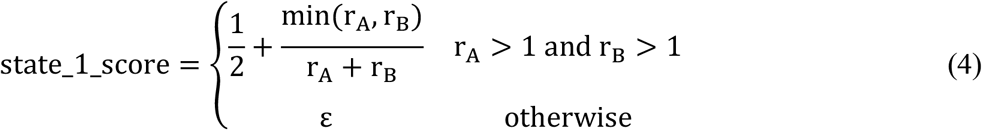

For all four types of relations above, state_0_score is complementary to state_1_score with respect to 1. For “*otherwise*“ condition in each equation, we set up a uniform parameter *ε* as the estimation for edge score. For each subroutine, there are 2 different sum scores corresponding to each binary sequence as previous defined. The binary subroutine occurrence probability (BSOP) averages along all simple paths between any arbitrary header-terminal protein pair for each binary sequence. For a subroutine with L edges, its mathematical form is given below:

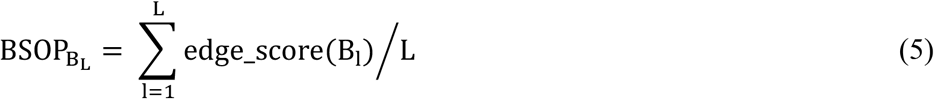

Here, *B_l_* denotes the binary input for the lth edge (PPI) and edge score is computed according to Equations (2) – (4) according to PPI relation with proper condition matching. All complete PPI subroutines with either state 0 or 1 occurrence probabilities greater than acceptable thresholds are merged to form the final disease network. The detailed description about loop sub-structure extraction has been described in [15].

### Post-treatment loop extraction model (dynamic model)

The dynamic evolutions of the loop sub-structure are studied with implementation of the following HMM algorithm. Discrete binary observations O are defined for each type of PPIs as follows. For activation/expression, binding/association and dissociation types of PPIs, we have:

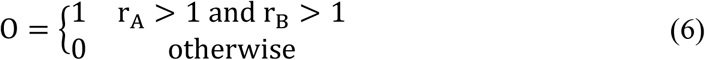

And for inhibition/repression type of PPIs, we have:

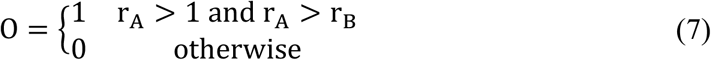

In this work, we propose a newly designed HMM initialization scheme as described in detail below that first generates time-series “fake” state sequences, which are partial outputs from the static model, and then the learned “supervised” HMM parameter pair from the “fake” sequence is selected as the initial parameter pair for the real unsupervised HMM problem. Finally, the Baum-Welch algorithm is used for iteration updates until the likelihood reaches its maximum. For our specific case, since HMM algorithms will be applied to study the dynamic behavior for each PPI, we therefore have a learned optimal triple pair *λ*\* = (*π*\*, *A*\*, *B*\*) with dimensions 2 by 1, 2 by 2 and 2 by 2 for each PPI. In the dynamic model, we utilize the probabilities inside the state transition matrix A as newly defined edge scores, so for each given PPI A to B, we update two edge scores (state_1_score and state_0_score) as:

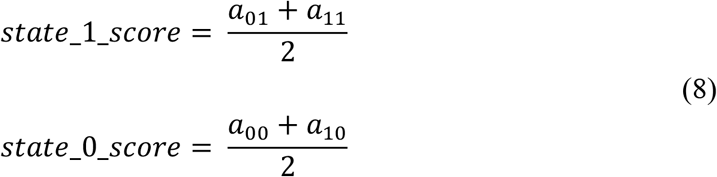

##### HMM with the newly designed initialization scheme

**Step 0: Generate multiple time-series “fake” state-observation sequences**

For each time point t = 1, 2, …, T:

> The discrete state sequences for difference types of PPI are defined according to Equations (2) – (4). Specifically, if the state_1_score for each type of PPI is > 0.5, the state is 1, otherwise is 0.

**Step 1: Generate initial HMM parameter set**

With the generated “fake” state-observation sequences, compute each parameter by counting the frequency, for example, components *a*_00_in the state transition matrix A, *π*_0_ in initial state vector, *b*_00_ in observation matrix can be estimated as follows:

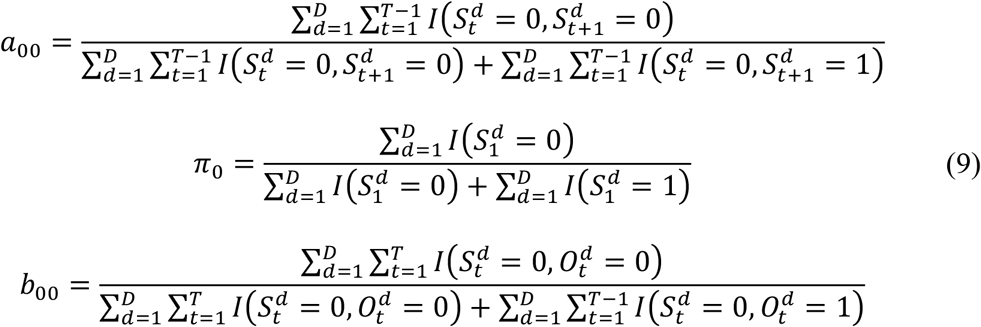

All rest components unlisted can be estimated similarly. In the equation, 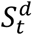 and 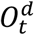 denote the state and observation of sample d at time t. In this way, we get the initial parameter set *λ*^(0)^ = (*π*^(0)^, *A*^(0)^, *B*^(0)^). Here *I* is the indicator function.

**Step 2: Iterative steps**

For k = 1, 2, …

Update the parameter set by Baum-Welch algorithm, and at each iterative step, with the current parameter set *λ*^(*k*)^ = (*π*^(*k*)^, *A*^(*k*)^, *B*^(*k*)^), compute the objective likelihood 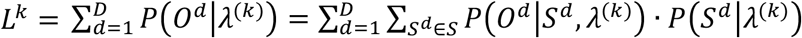. To be consistent with previous notations, the superscript d denotes sample index, therefore, both *O^d^* and *S^d^* are vectors of length T. In detail, we have:

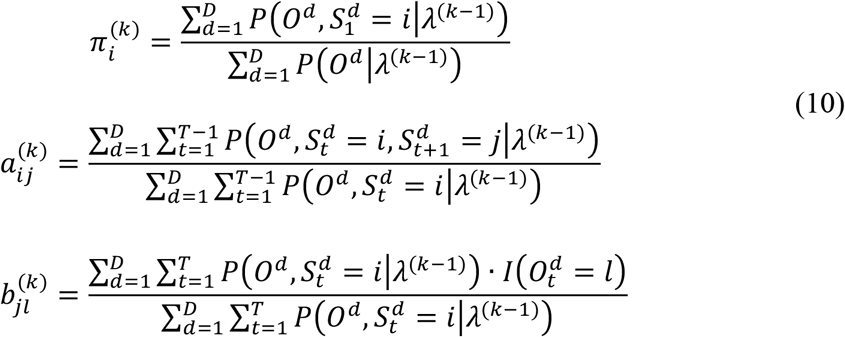

With

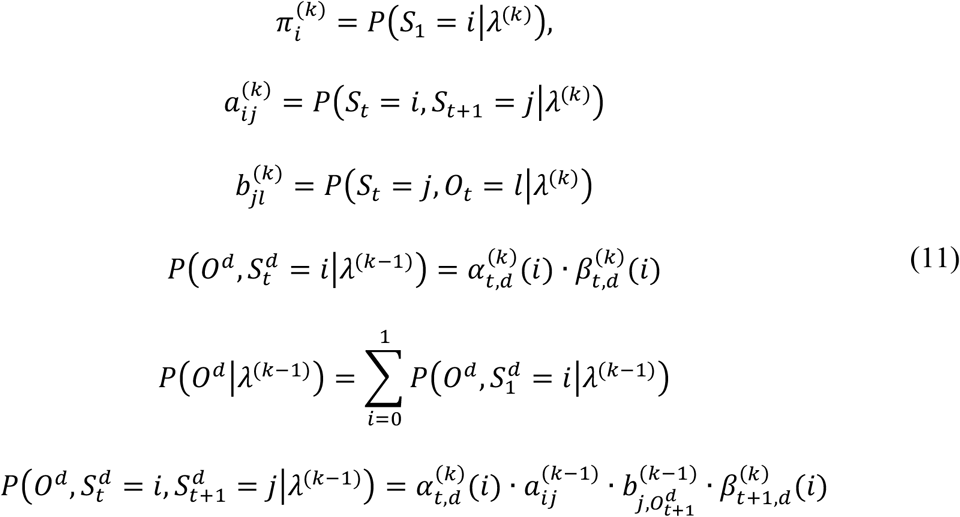

Here, 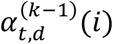 and 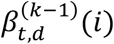 are updated recursively with forward and backward direction, separately.
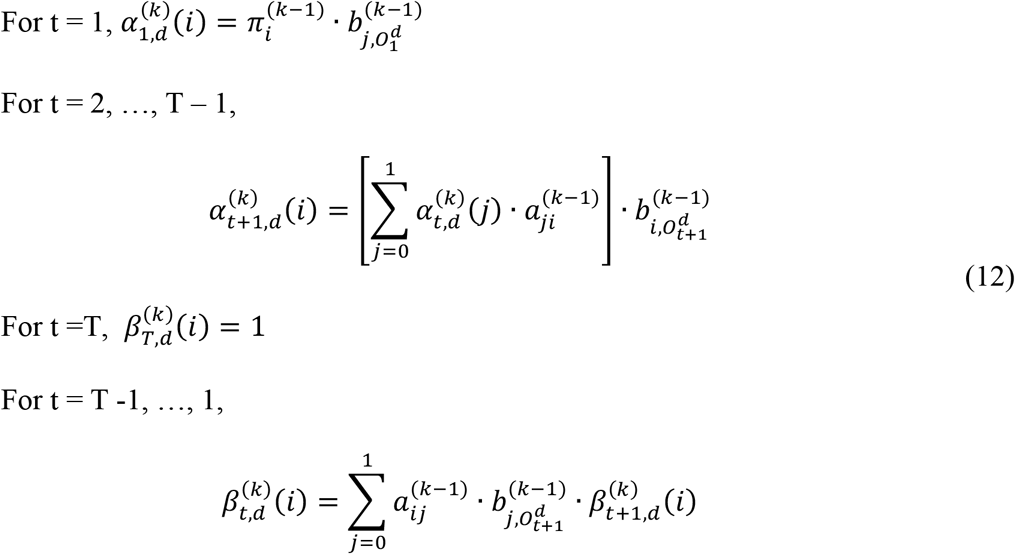

**Step 3: Stop the iteration**

Given *ε*, stop the iteration, when |*L*^*k*+1^ − *L^k^*| < *ε*.

### Evaluation of loop inhibition / activation ability for both static and dynamic models

The capability of the loop to inhibit / activate any protein X on the loop is computed in the following steps. Step 1: given any gene X from the loop, for any gene Y from the rest of the loop, find all shortest paths from genes Y to X and create corresponding two binary sequences for each path. Step 2: replace 0 and 1 in each binary sequence with the corresponding state_0_score and state_1_score. Step 3: for each shortest path, we can evaluate its failure (fails to inhibit gene X) and success (succeeds to inhibit gene X) scores by averaging the corresponding binary sequences separately. Step 4: for the whole loop impact to protein X, we repeat steps 1 – 3 for each of the remaining proteins on loop and collect all shortest paths with strong inhibition capabilities (average score > 0.95). We use the sum of all above threshold scores as the inhibition ability of the loop to protein X, applying a similar procedure to measure loop activation ability to protein X.

## Results

### Core signaling networks and loops of 3 MB subtypes (WNT, SHH and Group 3)

We used 2 signaling subroutine thresholds, 0.9 and 0.8, to obtain 3 signaling networks for each subtype separately. The extracted 3 loops are presented in **Fig. 1A, 1B and 1C**; the static model demonstrates its capability to distinguish the extracted loop substructures for these 3 MB subtypes to some extent, and the corresponding Venn diagrams under the highest threshold 0.9 are compared (see **Fig. 1D**). More detailed selected loop edges under different thresholds are presented in **Supplementary Table S1. Table 1** provides the hit rates for literature reported essential genes and drug targets of MB subtypes, with more details in **Supplementary File S1** and **Supplementary Table S1**. Drug targets were obtained from the DrugBank database [21]. For SHH and Group 3 subtype MBs, over 87.5% and 75.6% of reported therapeutic targets come from the complete loop substructure L_c_, and with threshold 0.8, the loops predicted by the static model cover 100% and 82.4% of all validated therapeutic targets for each MB subtype, respectively. In order to avoid confusion, it is necessary to explain the complete loop sub-structure L_c_ which is obtained by extracting the complete loop sub-structure directly based on provided PPI knowledge from the KEGG pathway database without considering any genomics data. Moreover, many missed targets are head genes, e.g., WNT3A, or the tail genes, e.g., PARP.

**Figure 1:**
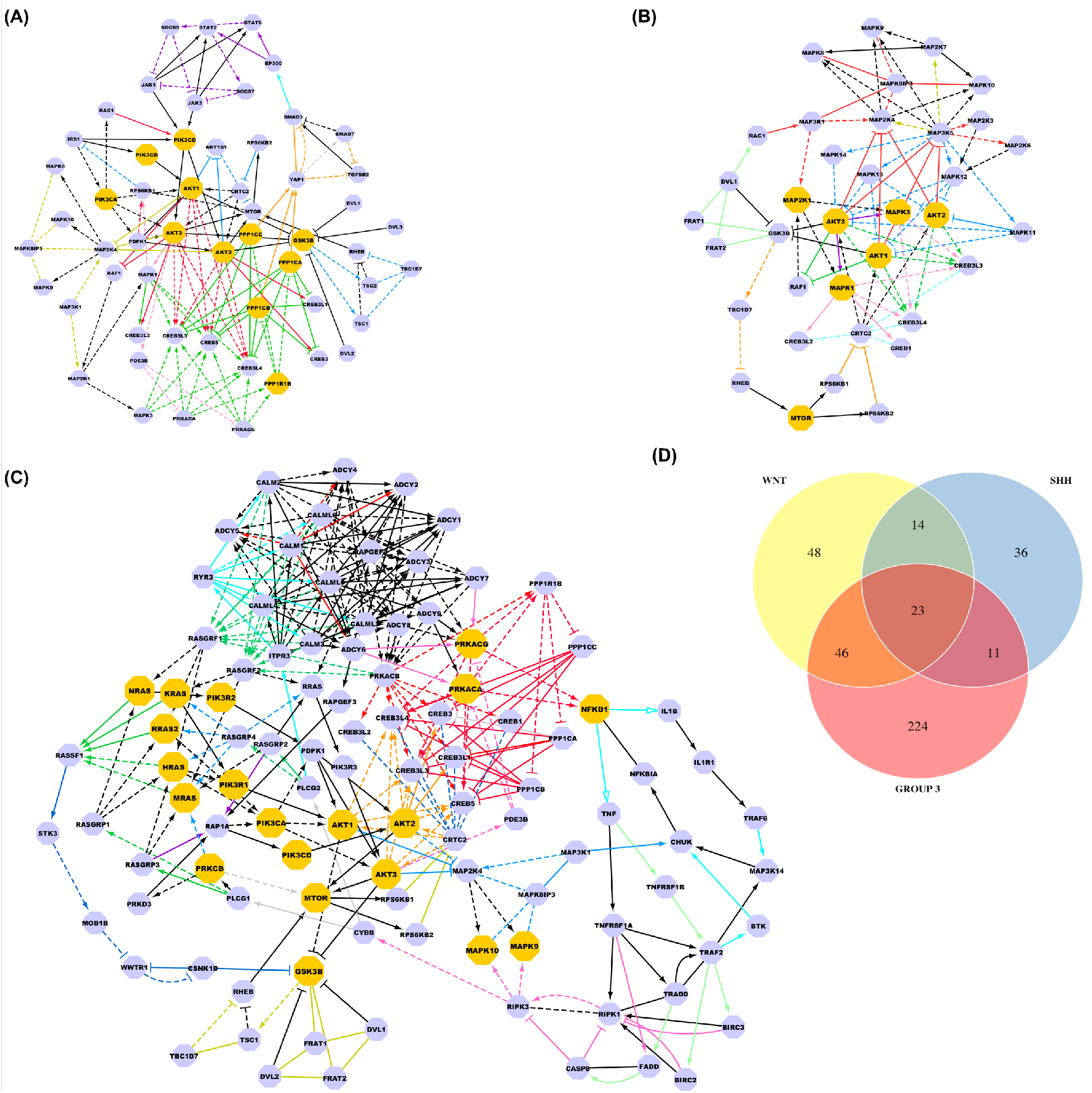
Larger yellow genes are reported therapeutic targets. Signaling interactions from different pathways are indicated with different colors. Dashed and solid lines denote the failure and successful occurrence of the provide PPIs, respectively. **(A):** Core signaling network and signaling loops of WNT MB. Color schemes: blue (MTOR only), red (PI3K-AKT only), green (CAMP only), purple (JAK-STAT only), orange (HIPPO only), pink (APELIN only), light blue (TGF-BETA only), dark yellow (MAPK only), and black (shared by multiple pathways). **(B):** Core signaling network and signaling loops of SHH MB. Color schemes: blue (Sphingolipid only), red (MAPK only), green (PI3K-AKT only), purple (APELIN only), orange (MTOR only), pink (CAMP only), light blue (AMPK only), dark yellow (TNF only), and black (shared by multiple pathways). **(C):** Core signaling network and signaling loops of Group 3 MB. Color schemes: blue (MAPK only), red (CAMP only), green (RAS only), purple (RAP-1 only), orange (PI3K-AKT only), pink (APELIN, Apoptosis, and Necroptosis only), light blue (Phospholipase D, NF-kappa_B, and CALCIUM), dark blue (AMPK and HIPPO), light green (TNF only), grey (HIF-1 only), dark yellow (MTOR and WNT only), and black (shared by multiple pathways). **(D):** The Venn diagram for 3 MB subtypes’ loop edges with highest subroutine threshold 0.9.

**Table 1:** Hit rates for literature reporting therapeutic targets for each of the 3 MB subtypes. The more detailed drug target information for each MB subtype is listed in **Supplementary Table S2**. The detailed literature references are attached in **Supplementary File S2**.

| **WNT type MBs** | **threshold = 0.9** | **threshold = 0.8** |
| --- | --- | --- |
| Numbers of reported therapeutic targets (total) | 10 | 10 |
| Numbers of reported therapeutic targets (from loop $L_c$) | 4 | 4 |
| Targets selected by algorithm (from loop $L_{0.8,0.9}$) | 3 | 4 |
| Hit rate | 75% | 100% |
| **SHH type MBs** | **threshold = 0.9** | **threshold = 0.8** |
| Numbers of reported therapeutic targets (total) | 16 | 16 |
| Numbers of reported therapeutic targets (from loop $L_c$) | 14 | 14 |
| Targets selected by algorithm (from loop $L_{0.8,0.9}$) | 4 | 14 |
| Hit rate | 28.6% | 100% |
| **Group 3 type MBs** | **threshold = 0.9** | **threshold = 0.8** |
| Numbers of reported therapeutic targets (total) | 45 | 45 |
| Numbers of reported therapeutic targets (from loop $L_c$) | 34 | 34 |
| Targets selected by algorithm (from loop $L_{0.8,0.9}$) | 12 | 28 |
| Hit rate | 35.3% | 82.4% |

### Numerical evaluation of dynamic model outputs

Two types of comparisons are made between the newly designed initialization scheme and random initialization: the maximum likelihoods computed and the cost of computational time. 50 random initializations are applied to each PPI. The computed likelihoods and cost computational time under 3 different schemes are attached in **Supplementary Table S2**. Here, we present the computed normalized likelihoods with different initialization schemes with respect to all activation PPI relations considered (see **Fig. 2**), the other 3 types of PPI relations are detailed in **Supplementary Fig. S1–3**. The complete learned state transition matrices for both datasets are attached in **Supplementary Table S3**. Sub-graphs in **Fig. 2** and **Supplementary Fig. S1–3** demonstrate that for most provided PPIs with 4 classified relations, the new initialization scheme can finally achieve comparable maximum likelihood when compared with random initializations. The robustness of the new initialization scheme is actually attributed to the rationality in the static model. As shown in **Fig. 2**, there are many red dots that could coincide with or are very close to green lines (constant horizontal lines with value 1.0), which means that for some PPIs, the computed likelihoods after one iteration with the newly designed initialization scheme are already quite close to their global maximum likelihoods. Specifically (See **Supplementary Table S2**), for activation type PPIs, 70.7% (GSE22875), 74.4% (GSE77042 with dose = 40 mg/kg), and 73.9% (GSE77042 with dose = 80 mg/kg) of the normalized likelihood ratios between one and multiple iterations are greater than 0.9 (rounded to the hundredth), similarly, for inhibition type PPIs, the corresponding ratios are 68.5% (GSE22875), 65.2% (GSE77042 with dose = 40 mg/kg), and 66.4% (GSE77042 with dose = 80 mg/kg), for binding/association type PPIs, the corresponding ratios are 73.1% (GSE22875), 74.9% (GSE77042 with dose = 40 mg/kg), and 76.8% (GSE77042 with dose = 80 mg/kg), for dissociation type PPIs, the corresponding ratios are 68.2% (GSE22875), 90.9% (GSE77042 with dose = 40 mg/kg) and 81.8% (GSE77042 with dose = 80 mg/kg). This phenomenon from the other perspective, demonstrates the rationality in the static model, in detail, e.g., for activation PPIs, by considering RNA expression alone, when *r_A_* > 1 and *r_B_* > 1, and the state_1_score is greater than 0.5, then it is quite likely that protein A could activate protein B in real.

**Figure 2:**
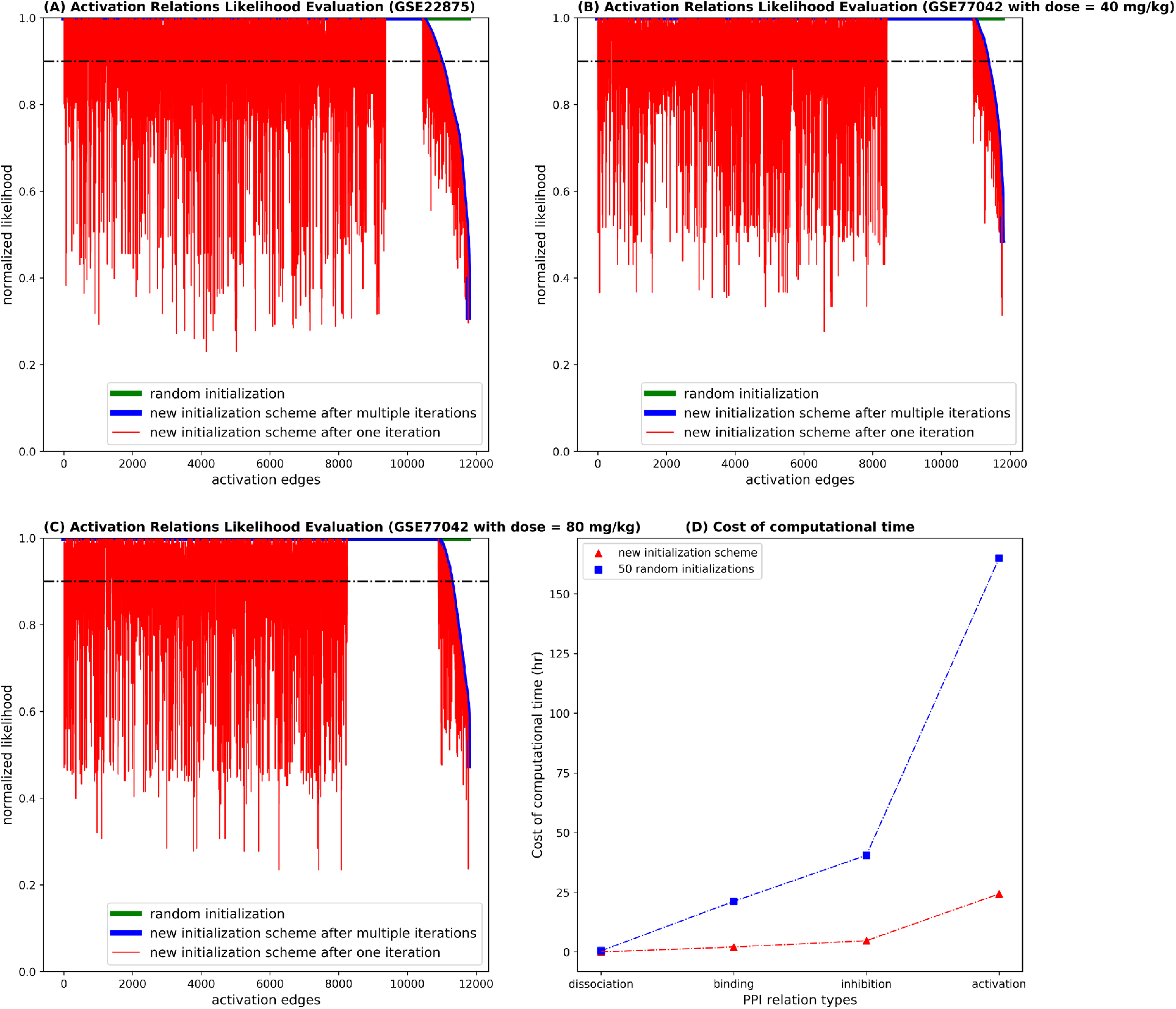
The comparisons of computed likelihoods with different initialization schemes for all activation PPI types of relations. Among the total 11774 activation edges considered, for **(A):** dataset GSE22875, dataset GSE77042 with **(B):** dose 40 mg/kg and with **(C):** dose 80 mg/kg, the computed maximum likelihoods of 10976 (93.2%), 11336 (96.3%), and 11272 (95.7%) activation edges by the new initialization scheme have the normalized likelihood (= computed_likelihood_by_scheme X/ computed_likelihood_by_random_initialization) > 0.9. Scheme X could be new initialization scheme, and new initialization scheme after one iteration. **(D):** comparison of computational times is presented, and for each type of PPI, the new initialization scheme costs only 1/7 of the time that it would take if starting with 50 random initializations.

### Biological evaluation of dynamic model outputs — Increased upstream inhibition signaling (PI3K-AKT, PI3K-AKT themselves are inhibited) after OTX2 silencing

Based on KEGG pathway database, there are different subroutines in PI3K-AKT and TGF-*β* signaling transduction pathways that finally inhibit MYC. In this sub-subsection, we first determine if any subroutine fails to inhibit MYC (state_0_score > 50%) in dynamic manner after treatment. It is interesting to note that, according to our prediction results, there are 3019 (around 9.6% of the total) different subroutines in PI3K-AKT that will fail to inhibit MYC after treatment. Next, we are interested in how many subroutines in PI3K-AKT pathway could successfully inhibit MYC (state_1_score > 50%). Based on our prediction results, there are 28087 (around 89.1% of the total, 2.18 times of that before treatment) different subroutines that could inhibit MYC after treatment, 28018 of which can successfully inhibit MYC by inhibiting PI3K-AKT protein family that has been validated on exactly the same cell line (direct validation of up-regulation of PTEN) after reducing OTX2 level [22]. In the same work, it is also reported that OTX2 level is positively correlated with TGF-*β* signaling activity. Therefore, TGF-*β* signaling pathway should NOT have the capability to inhibit MYC after OTX2 silencing, and our results identify 47 (shrinks to 38.5% of that before treatment) subroutines from TGF-*β* pathway which could numerically inhibit MYC after treatment. All the computational details about these 2 pathways are presented in **Supplementary Table S4**. The detailed pre- and post-treatment comparisons are described in **Table 2**.

**Table 2:**
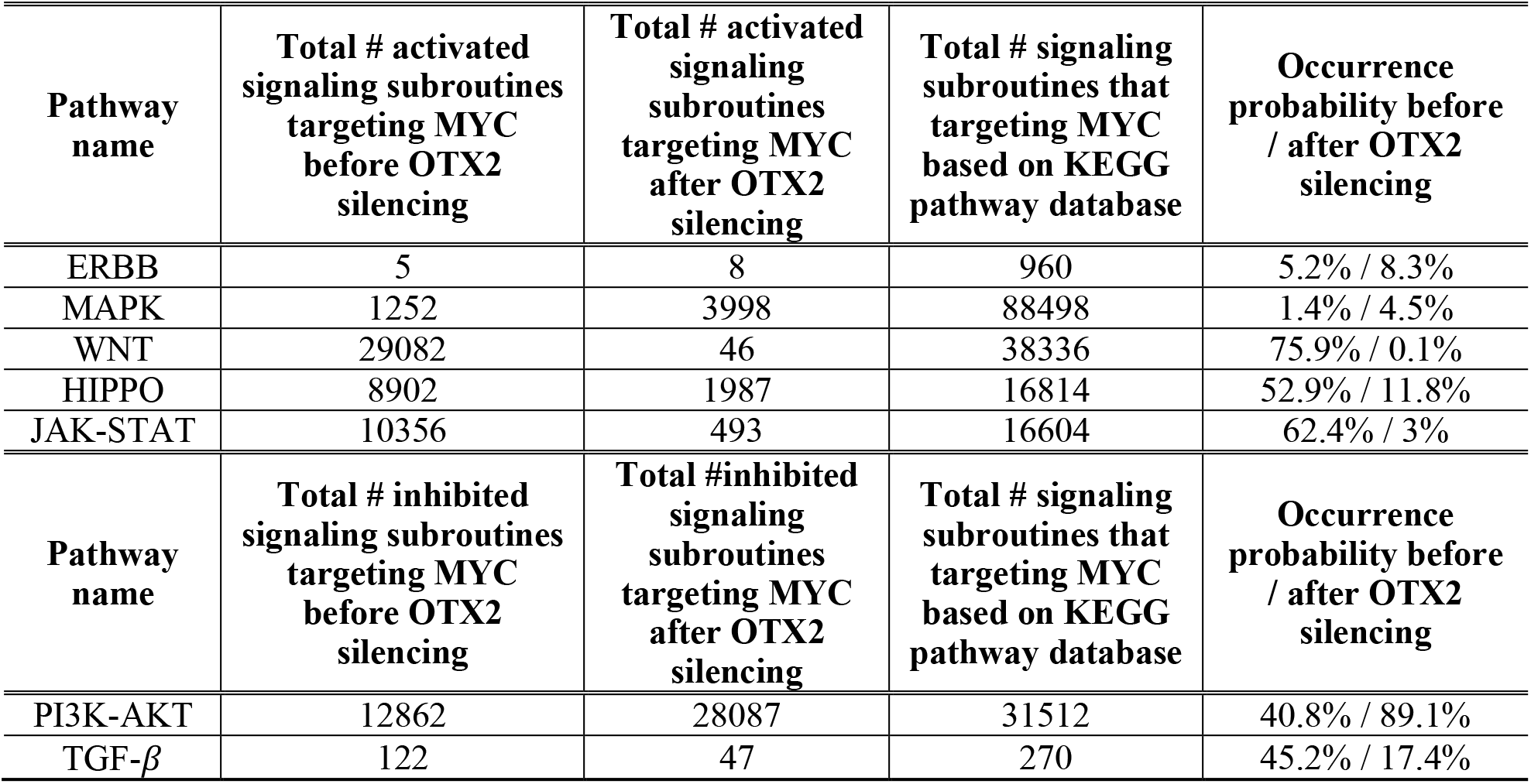
Occurrence probabilities for those subroutines that either activate or inhibit MYC in different signaling transduction pathways according to KEGG pathway database before and after OTX2 inhibition.

| Pathway name | Total # activated signaling subroutines targeting MYC before OTX2 silencing | Total # activated signaling subroutines targeting MYC after OTX2 silencing | Total # signaling subroutines that targeting MYC based on KEGG pathway database | Occurrence probability before / after OTX2 silencing |
| --- | --- | --- | --- | --- |
| ERBB | 5 | 8 | 960 | 5.2% / 8.3% |
| MAPK | 1252 | 3998 | 88498 | 1.4% / 4.5% |
| WNT | 29082 | 46 | 38336 | 75.9% / 0.1% |
| HIPPO | 8902 | 1987 | 16814 | 52.9% / 11.8% |
| JAK-STAT | 10356 | 493 | 16604 | 62.4% / 3% |
| Pathway name | Total # inhibited signaling subroutines targeting MYC before OTX2 silencing | Total #inhibited signaling subroutines targeting MYC after OTX2 silencing | Total # signaling subroutines that targeting MYC based on KEGG pathway database | Occurrence probability before / after OTX2 silencing |
| PI3K-AKT | 12862 | 28087 | 31512 | 40.8% / 89.1% |
| TGF-β | 122 | 47 | 270 | 45.2% / 17.4% |

### Biological evaluation of dynamic model outputs — Decreased upstream activation signaling (WNT, HIPPO and JAK-STAT) after OTX2 inhibition

Even though the behaviors of subroutines that could activate MYC are not mentioned [22], we still investigate the simulated behaviors of those subroutines after OTX2 silencing in our study. MYC could be finally activated by different subroutines in ERBB, MAPK, WNT, HIPP, and JAK-STAT signaling transduction pathways in the KEGG pathway database. In this sub-subsection, we will determine if all subroutines targeting the oncogene MYC in different signaling pathways are de-activated in time-dependent manner after OTX2 silencing. The evaluation criterion is for those subroutines that finally activate MYC, we check their occurrence frequencies in the activated states (state_1_score > 50%), and the occurrence probabilities before and after OTX2 silencing are summarized in **Table 2** below. All aforementioned signaling pathways that could activate MYC show very low activation probabilities after OTX2 silencing (< 12%), therefore, it is fair to say those signaling transduction pathways are de-activated after OTX2 silencing. All scored subroutines that finally activated MYC are attached in **Supplementary Table S4**.

### Hypothesize loop as the measurement of drug resistance evaluation and prediction

In previous sub-sections, post-OTX2 silencing behaviors of different signaling pathways that could either activate or inhibit MYC are studied independently, and it is quite obvious that after OTX2 silencing, WNT, JAK-STAT, and HIPPO signaling pathways are strongly inhibited, but the MAPK pathway is more highly activated. This phenomenon shows that after effective treatments, some pathways which positively contributed to tumor development can definitely be suppressed, however, other similarly functioned (e.g. activating MYC) pathways could become even more activated. This motivates us to boldly hypothesize that the loop sub-structure which considers the crossing talking of various pathways concurrently should be considered as a reasonable explanation for drug resistance. Under our hypothesis, a treatment (drug / drug combo) is efficient when after treatment, the loop demonstrates stronger inhibition abilities to cell cycle and DNA replication related proteins. To clarify, when we apply loop to explain the drug resistance, we use the complete loop extracted from all KEGG pathways considered, not just the before-treatment partial disease loop.

### Effective treatment validates loop hypothesis: except CDK1, all rest cell cycle proteins get inhibited after considering cross talk of multiple pathways

In the paper [23] (dataset GSE22875), different gene ontology (GO) [24] annotated gene sets related to cell cycle and DNA regulations are proven to be strongly down-regulated after silencing OTX2. Therefore, we first prove that biologically validated down-regulated genes in dataset GSE22875 which also sat on the loop can be validated numerically for stable down-regulation by considering the cross talk of multiple pathways and post-treatment time-series data. Next, we compare numbers of strong inhibition and activation shortest paths for the same set of cell cycle and DNA replication proteins between pre- and post-treatment conditions for Group 3 type MBs under the loop hypothesis. The complete considered GO annotated gene sets and KEGG annotated cell cycle check points (only intersections with pathways mentioned ahead), and their topological distribution among the disease network are summarized in **Supplementary Table S5**. The computed loop inhibition and activation capabilities are presented in **Supplementary Table S5**. For dataset GSE22875, among the 33 cell cycle and DNA replication related proteins, mRNA expression level of CCND3, CDK2, E2F2/3 and MYC have been biologically validated to be strongly down-regulated, as are the protein levels of CCND3 and MYC [23]. When compared to the numerical simulation, all above mentioned proteins (100% agreements) are predicted to be consistently inhibited according to the mathematical assumptions embedded within the HMM algorithm. For one group 3 MB driver mutation gene MYC, when we were validating inhibition of MYC by considering individual KEGG pathways independently, it is interesting to note that there are still many different subroutines that have higher probabilities to activate MYC, but when finally merging into the loop structure which covers the cross talks between different signaling pathways presents much stronger global inhibition (948 V.S. 34) capability. Both strong successful inhibition and unsuccessful inhibition shortest paths in loop targeting MYC are presented in **Supplementary Fig. S4**. All strongly MYC successful and unsuccessful post-treatment inhibition subroutines (break into edges) are attached in **Supplementary Table S6**. As aforementioned, after OTX2 silencing, the inhibited PI3K-AKT pathway activity reports its indirect (directly related to PTEN) contribution to tumor suppression. Therefore, the other exciting find is that even under really high average threshold 0.95, multiple PPIs from the PI3K-AKT pathway do appear as strongly successful inhibition subroutines.

To further illustrate the loop hypothesis, comparison analyses analogous to **Table 2** are performed for the same set of cell cycle and DNA replication proteins by considering the integrative cross talk of various signaling transduction and cellular process pathways under both before and after treatment conditions. Specifically, **Table 3** presents numbers of strong inhibition and activation shortest paths targeting representative cell cycle phase check points for both before and after treatments for comparison. The complete comparisons for all GO and KEGG annotated cell cycle and DNA replication proteins are listed in **Supplementary Table S5**.

**Table 3:** Comparison of numbers of strong (average path score > 0.95) inhibition and activation shortest paths for representative cell cycle phase check points between before and after treatments conditions for Group 3 subtype MBs under the loop hypothesis (considering cross talk of multiple pathways instead).

| **# of Inhibition shortest paths / # of Activation shortest paths (Group 3 MBs)** |  |  |
| --- | --- | --- |
| **Gene Symbol** | **Before Treatment** | **After Treatment** |
| MYC | 2 / 517 | 948 / 34 |
| CDK1 | 47 / 2787 | 138 / 2028 |
| CDK2 | 549 / 248 | 993 / 2 |
| CDK4 | 0 / 3206 | 374 / 49 |
| CDK6 | 150 / 0 | 3 / 2 |
| CCNA1 | 0 / 666 | 158 / 0 |
| CCNA2 | 0 / 1331 | 597 / 0 |
| CCNB1 | 1925 / 5 | 355 / 129 |
| CCNB2 | 1925 / 4 | 355 / 197 |
| CCND1 | 810 / 246 | 1573 / 37 |
| CCND2 | 732 / 693 | 455 / 126 |
| CCND3 | 624 / 0 | 977 / 35 |
| CCNE1 | 537 / 572 | 1982 / 0 |
| CCNE2 | 0 / 881 | 599 / 3 |
| E2F2 | 1013 / 0 | 1376 / 218 |
| E2F3 | 0 / 3143 | 1381 / 216 |

### Ineffective treatments validate loop hypothesis: most cell cycle proteins fail to be inhibited after considering cross talk of multiple pathways

The computed dynamic PPI probabilities for GSE77042 are based on treatments with doses = 40 and 80 mg/kg once a day. According to biological experiments, MK-4101 treatments with doses of both 40 and 80 mg/kg each day failed to inhibit tumor growth (see **Fig. 1B and C** in [25]). Gene expression analysis in **Fig. 5D** in [25] concluded the same: at doses = 40 and 80 mg/kg each day, cell cycle related genes do not present strongly stable inhibited evidence at all in time evolved manner. When comparing with simulation results, for genes related to G1-S transitions, the loop presents much higher activation capabilities than most representative proteins. The loop with dose = 80 mg/kg once a day illustrates much stronger inhibition impact compared that with dose 40 mg/kg once a day (most blue squares are above red triangles, see **Supplementary Fig. S5**). Similar trends have been shown in other biological experimental results, e.g., tumors treated with 80 mg/kg once a day present lower tumor volume growth rate and mice body weight change compared with those treated with 40 mg/kg once a day. Similarly, for dataset GSE77042, the comparison of numbers of strong inhibition and activation paths for representative cell cycle check points between before and after treatment conditions are presented in **Table 4**, and the more detailed information are attached in **Supplementary Table S5**.

**Table 4:** Comparison of numbers of strong (average path score > 0.95) inhibition and activation shortest paths for representative cell cycle phase check points between before and after treatments conditions for SHH subtype MBs under the loop hypothesis (considering cross talk of multiple pathways instead).

| **# of Inhibition shortest paths / # of Activation shortest paths (SHH MBs)** |  |  |  |
| --- | --- | --- | --- |
| **Gene Symbol** | **Before Treatment** | **After Treatment (dose = 40 mg/kg)** | **After Treatment (dose = 80 mg/kg)** |
| CDK1 | 107 / 530 | 0 / 17 | 0 / 1593 |
| CDK2 | 0 / 455 | 57 / 1544 | 0 / 0 |
| CDK4 | 0 / 1928 | 85 / 2074 | 99 / 164 |
| CDK6 | 0 / 1915 | 86 / 1552 | 99 / 159 |
| CCNA1 | 0 / 215 | 61 / 985 | 52 / 1 |
| CCNA2 | 0 / 401 | 370 / 1190 | 52 / 0 |
| CCNB1 | 409 / 28 | 0 / 0 | 0 / 22 |
| CCNB2 | 409 / 4 | 1491 / 1 | 767 / 22 |
| CCND1 | 222 / 103 | 1397 / 253 | 1130 / 58 |
| CCND2 | 216 / 537 | 1072 / 324 | 616 / 125 |
| CCND3 | 49 / 3 | 1002 / 1725 | 361 / 87 |
| CCNE1 | 0 / 521 | 608 / 1542 | 789 / 0 |
| CCNE2 | 0 / 507 | 375 / 1208 | 14 / 0 |
| E2F2 | 220 / 0 | 1 / 1980 | 0 / 33 |
| E2F3 | 0 / 1292 | 177 / 1980 | 0 / 33 |

For effective treatments (see **Table 3**), except CDK1, the post-treatment loop presents much stronger inhibition capabilities to all the rest cell cycle check point proteins compared to activation capabilities. For most cell cycle check point proteins, sharp contrasts between inhibition and activation capabilities can be directly observed when comparing pre-treatment and posttreatment loops. However, for ineffective treatments (see **Table 4**), for most cell cycle check point proteins, there are no inhibition and activation capabilities differences between pre-treat and post-treat loops, and all three loops present much stronger activation capabilities. The loop state_0_score and state_1_score for both Group 3 and SHH MBs under pre- and post-treatment conditions are attached in **Supplementary Table S6** as well.

### Both pre- and post-treatment loops present unique set of strongly “activated” and “deactivated” loop edges

After validating and explaining the loop theory, the more interesting subsequent question is: what is the difference between pre- and post-treatment loops? This question will be addressed firstly by considering both strongly “activated” and “de-activated” (here, strongly “activated” here means state_1_score > 0.9, and strongly “de-activated” equals state_0_score > 0.9) edges within the same loop under pre and post-treatment conditions. Different comparison results with threshold = 0.9 are presented in **Supplementary Table S7**. The corresponding pie chart plot is attached as **Fig. 3A**. First, it is interesting to note that despite considering strongly “activated” or “de-activated” edges, both have around half (50%) of themselves exclusively, i.e., a unique set of strongly deregulated loop edges before treatment and simultaneously another unique set of strongly deregulated loop edges after treatment exist. This echoes with what we stressed before when validating the loop hypothesis: we should consider the full complete loop, not any other partial structures, since it is nonsensical to assume that after treatment, only the loop sub-structure which is closely related to the disease pre-treatment loop gets strongly influenced. Importantly, 41.8% of “activated” and 4.8% of “de-activated” pre-treatment loop edges switch to opposite original status after treatment, but 5.4% of “activated” and 36.7% of “deactivated” pre-treatment loop edges keep their original status. All these observations and cross verifications demonstrate that the loop theory should be considered as a very important tool capable of characterizing the drug effects and forecasting drug resistance.

**Figure 3:**
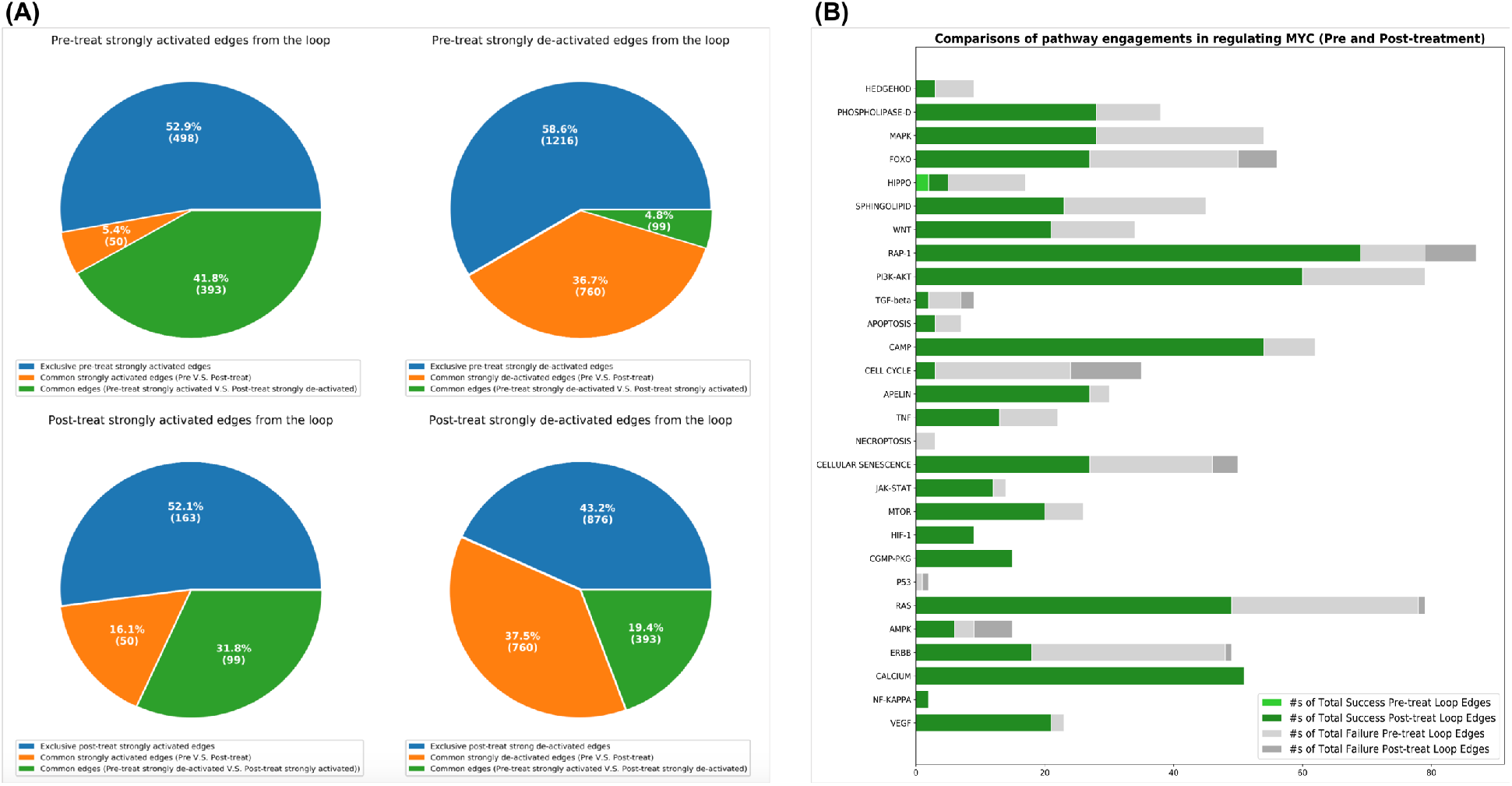
**(A):** Visualized comparisons between pre and post-treatment loops (threshold = 0.9). **(B):** Comparisons of pathway engagements in regulating MYC when considering both pre and post-treatment conditions. “Success” is defined as successful inhibition + unsuccessful activation and “Failure” is defined as successful activation + unsuccessful inhibition.

### Comparisons of pathway engagements under pre and post-treatment conditions: almost all pathways present much stronger capabilities in inhibiting all cell cycle related proteins

In this sub-section, comparison of pathway engagements in either activating or inhibiting cell cycle and DNA replication proteins under both pre- and post-treatment conditions will be evaluated, respectively. For each cell cycle and DNA replication related proteins, similar analyses with different comparisons will be performed individually for each pathway. Tabular analysis results are collected in **Supplementary Table S7** (sheets with format = “protein_STAT”), a more intuitive MYC graphical presentation is attached below (see **Fig. 3B**), and the remaining representative graphical representations are plotted in **Fig. 4A, 4B and 4C**. All cell cycle and DNA replication proteins’ successful and unsuccessful activation and inhibition loop edges with pathway annotations are attached in **Supplementary Table S7** (sheets with format = “protein_LOOP_EDGES”). Strikingly, for all cell cycle and DNA replication related proteins, almost all pathways present much stronger inhibition and much weaker activation capabilities after effective treatment.

**Figure 4:**
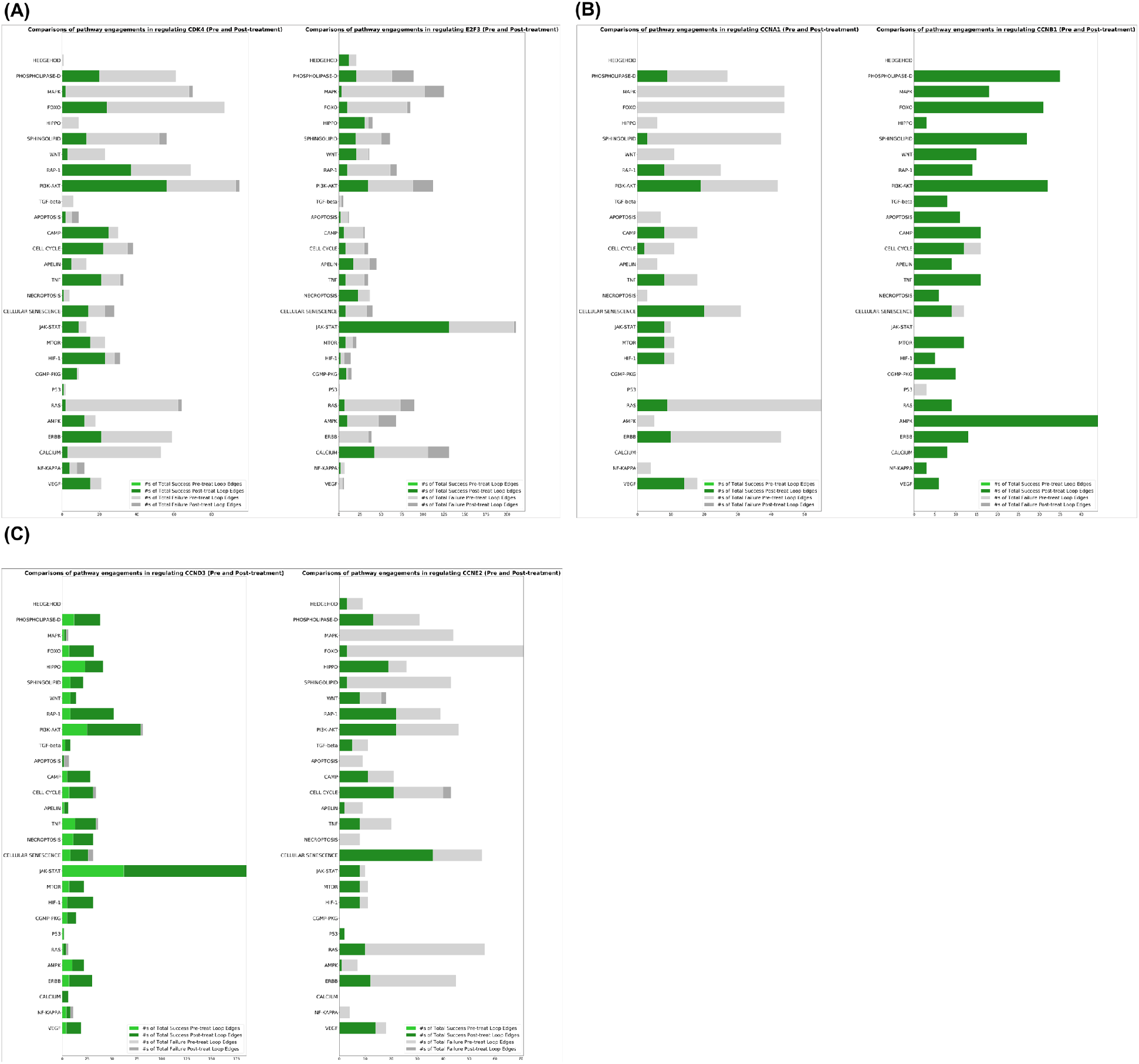
Comparisons of pathway engagements in regulating **(A):** CDK4 (left) and E2F3 (right), **(B):** CCNA1 (left) and CCNB1 (right), **(C):** CCND3 (left) and CCNE2 (right), when considering both pre and post-treatment conditions. “Success” is defined as successful inhibition + unsuccessful activation and “Failure” is defined as successful activation + unsuccessful inhibition.

## Discussion and Conclusions

In this paper, we hypothesize that the signaling loop sub-structures embedded in the PPI network in regulating driver mutation genes, cell cycle and DNA replication genes can be used as a tool to interpret and predict drug resistance. Both static and dynamic (HMM) models with newly designed initialization scheme which infer PPI static and dynamic activity probabilities separately are proposed to facilitate hypothesis validation and explanation. The HMM dynamic model with this newly designed initialization scheme presents promising stability and accurateness under both numerical and biological examinations and multiple datasets. This initialization scheme should work quite well for other datasets as well. The loop hypothesis is validated by multiple comparisons, i.e., comparisons between pre- and post-treatments, efficient and ineffective treatments. The signaling loop presents sharp contrasts (see **Table 3**) in regulating cell cycle and DNA replications proteins when compared between pre- and post-effective-treatments, however, when treating with ineffective options, the loop presents no apparent regulating capability difference for partial or most of the cell cycle and DNA replication proteins (see **Table 4**). Furthermore, with further investigation, it is very exciting to observe that almost all pathways within the loop play a larger role in controlling each cell cycle and DNA replication protein. Also, when comparing the pre and post-effective treatment loops, we find that partial previously highly activated loop edges are successfully de-activated after effective treatments, however, it is also shown that both pre and post-effective treatment loops have around half of their loop edges either highly activated or de-activated exclusively, and this provides rationale for working with the complete loop-substructure embedded in the PPI network, rather than just focusing on a partial loop which is only related to disease network (before treatments). Unfortunately, we must point out that the due to the limited availability of public medulloblastoma data, we could not find the RNA expression data for both group 3 medulloblastoma cell line D425 and the control normal cerebellum samples from the same platform. Similar embarrassment holds for dataset GSE77042 as well, so we use different patients’ samples instead as the input for the static model in order to present the comparison. In future work, the accuracy of the static model may improve with the incorporation multiple types of data (mutation, methylation, etc.), and the consideration of more pathways with other functionalities (e.g., metabolism). Lastly and most importantly, promising single drug and drug combination prediction algorithms which take drug resistance as one of the most important drug effect indicators will be developed based on the loop hypothesis in future.

## Supplementary Figures

**Supplementary Figure S1:**
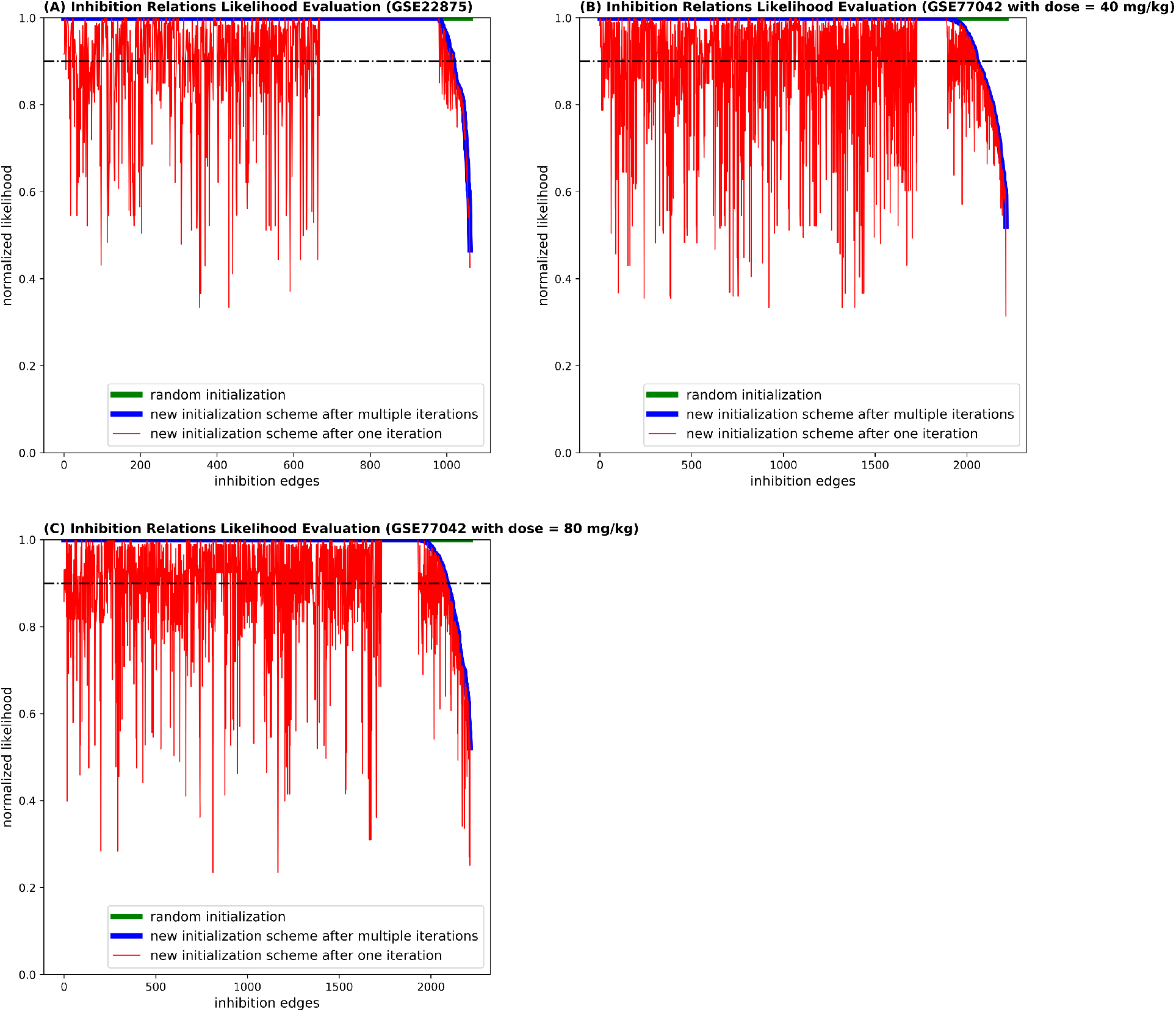
The comparisons of computed likelihoods with different initialization schemes for all inhibition PPI types of relations. Among the total 2214 inhibition edges considered, for (A): dataset GSE22875, dataset GSE77042 with (B): dose 40 mg/kg and with (C): dose 80 mg/kg, the computed maximum likelihoods of 2025 (91.5%), 2056 (92.9%), and 2087 (94.3%) inhibition edges by the new initialization scheme have the normalized likelihood (= computed_likelihood_by_scheme X/ computed_likelihood_by_random_initialization) > 0.9. Scheme X could be new initialization scheme, and new initialization after one iteration.

**Supplementary Figure S2:**
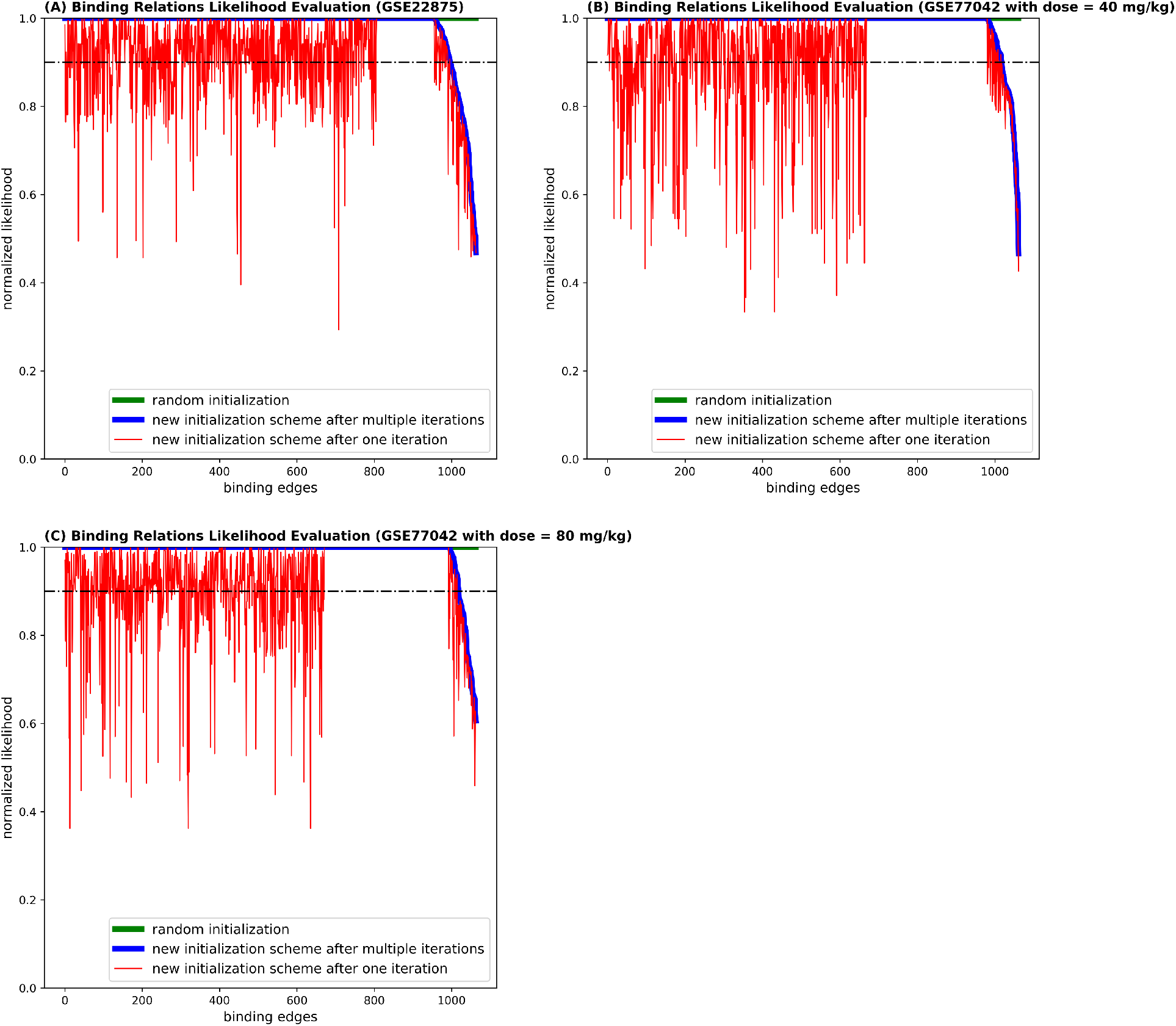
The comparisons of computed likelihoods with different initialization schemes for all binding PPI types of relations. Among the total 1064 binding edges considered, for (A): dataset GSE22875, dataset GSE77042 with (B): dose 40 mg/kg and with (C): dose 80 mg/kg, the computed maximum likelihoods of 997 (93.7%), 1018 (95.7%), and 1019 (95.8%) binding edges by the new initialization scheme have the normalized likelihood (= computed_likelihood_by_scheme X/ computed_likelihood_by_random_initialization) > 0.9. Scheme X could be new initialization scheme, and new initialization scheme after one iteration.

**Supplementary Figure S3:**
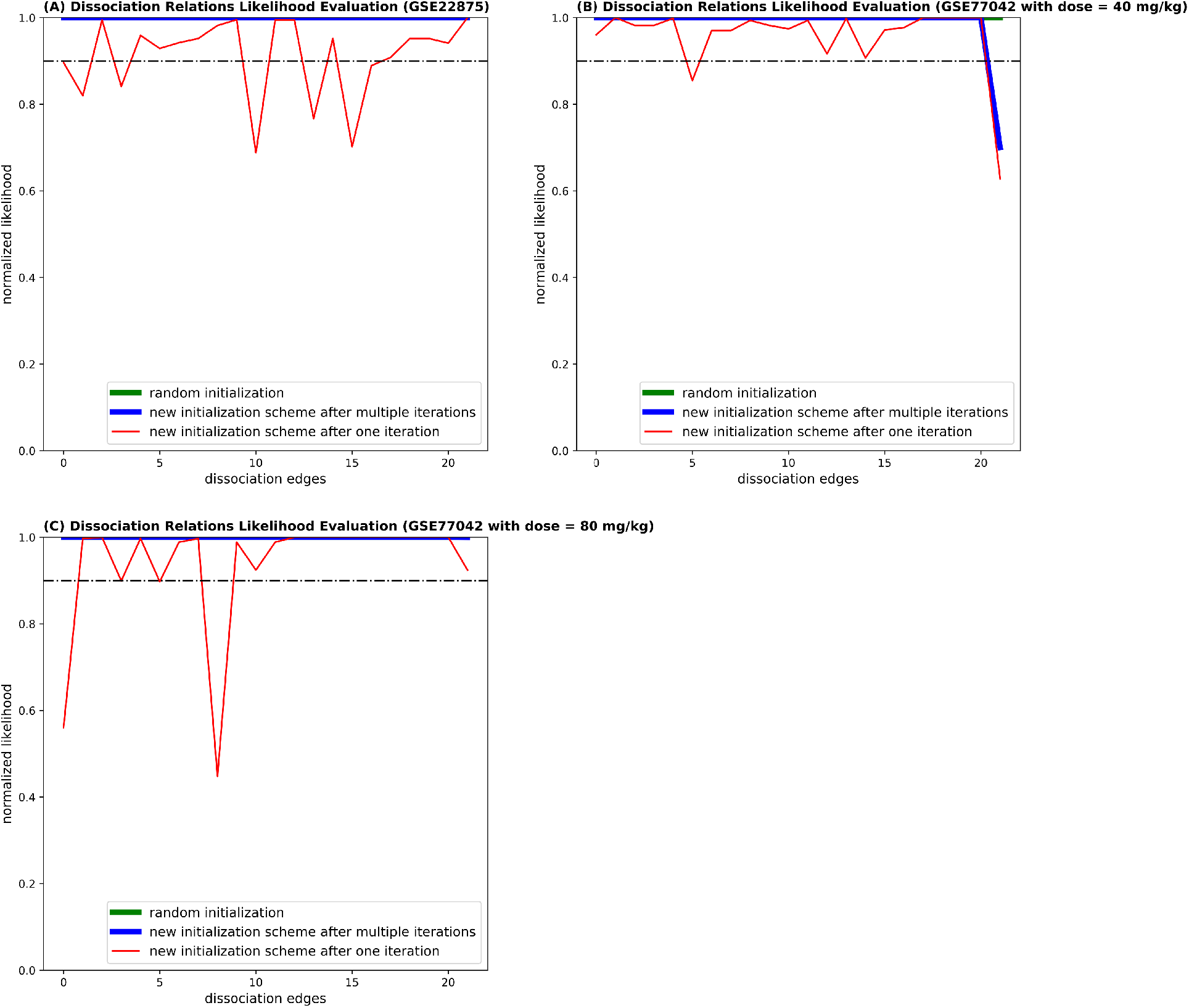
The comparisons of computed likelihoods with different initialization schemes for all dissociation PPI types of relations. Among the total 22 dissociation edges considered, for (A): dataset GSE22875, dataset GSE77042 with (B): dose 40 mg/kg and with (C): dose 80 mg/kg, the computed maximum likelihoods of 22 (100%), 21 (95.5%), and 22 (100%) dissociation edges by the new initialization scheme have the normalized likelihood (= computed_likelihood_by_scheme X/ computed_likelihood_by_random_initialization) > 0.9. Scheme X could be new initialization scheme, and new initialization scheme after one iteration.

**Supplementary Figure S4:**
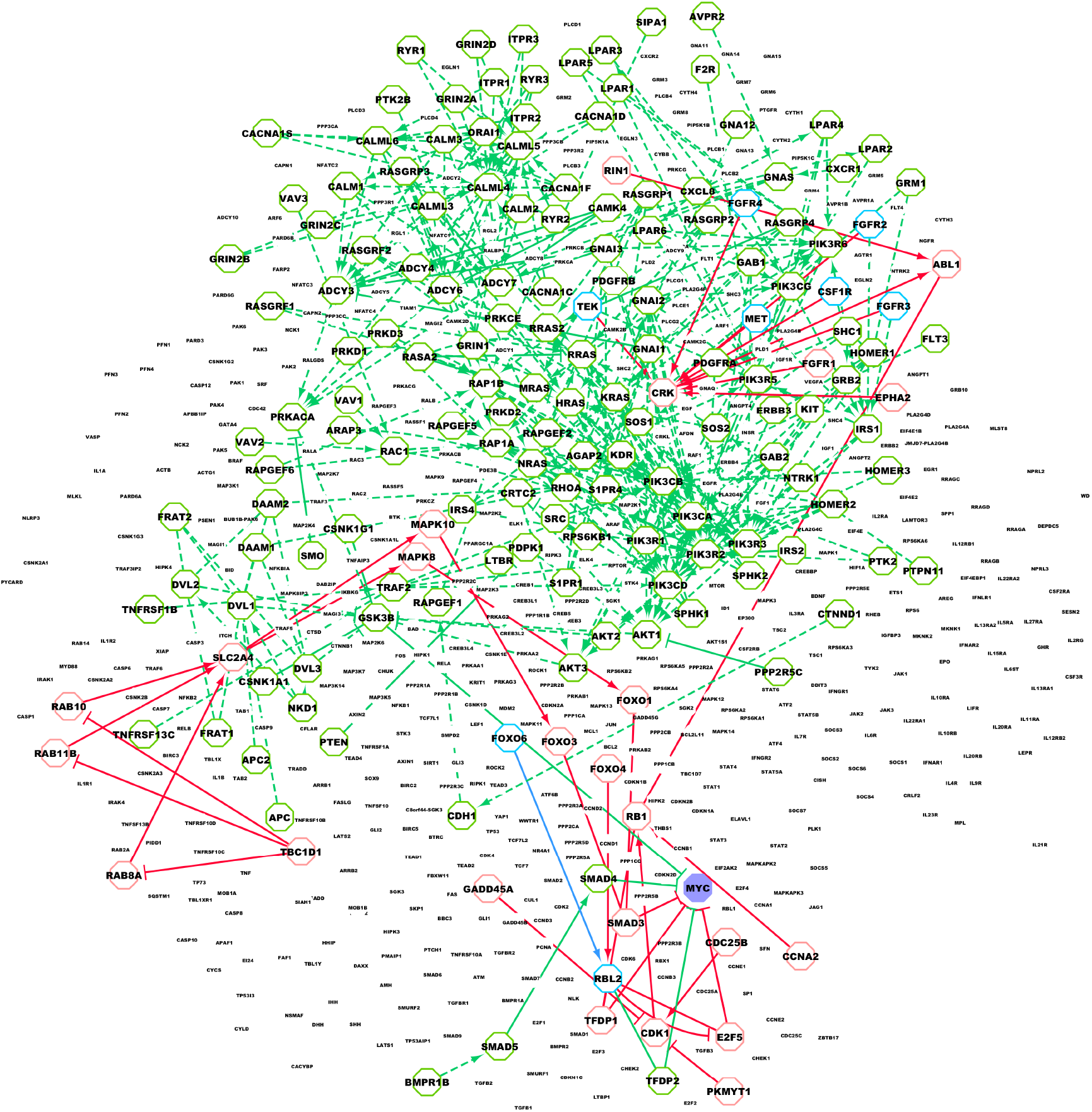
High (score > 0.95) inhibition (green) and activation (red) shortest paths to MYC. White background nodes are rest loop nodes. Enlarged purple node is gene MYC, enlarged light green boarded nodes are proteins that on paths which presents stronger inhibition capabilities to gene MYC only, enlarged light red boarded nodes are proteins that on paths which presents stronger activation capabilities to gene MYC only, and enlarged light blue boarded nodes are proteins that on paths which presents both strong inhibition and activation capabilities to gene MYC.

**Supplementary Figure S5:**
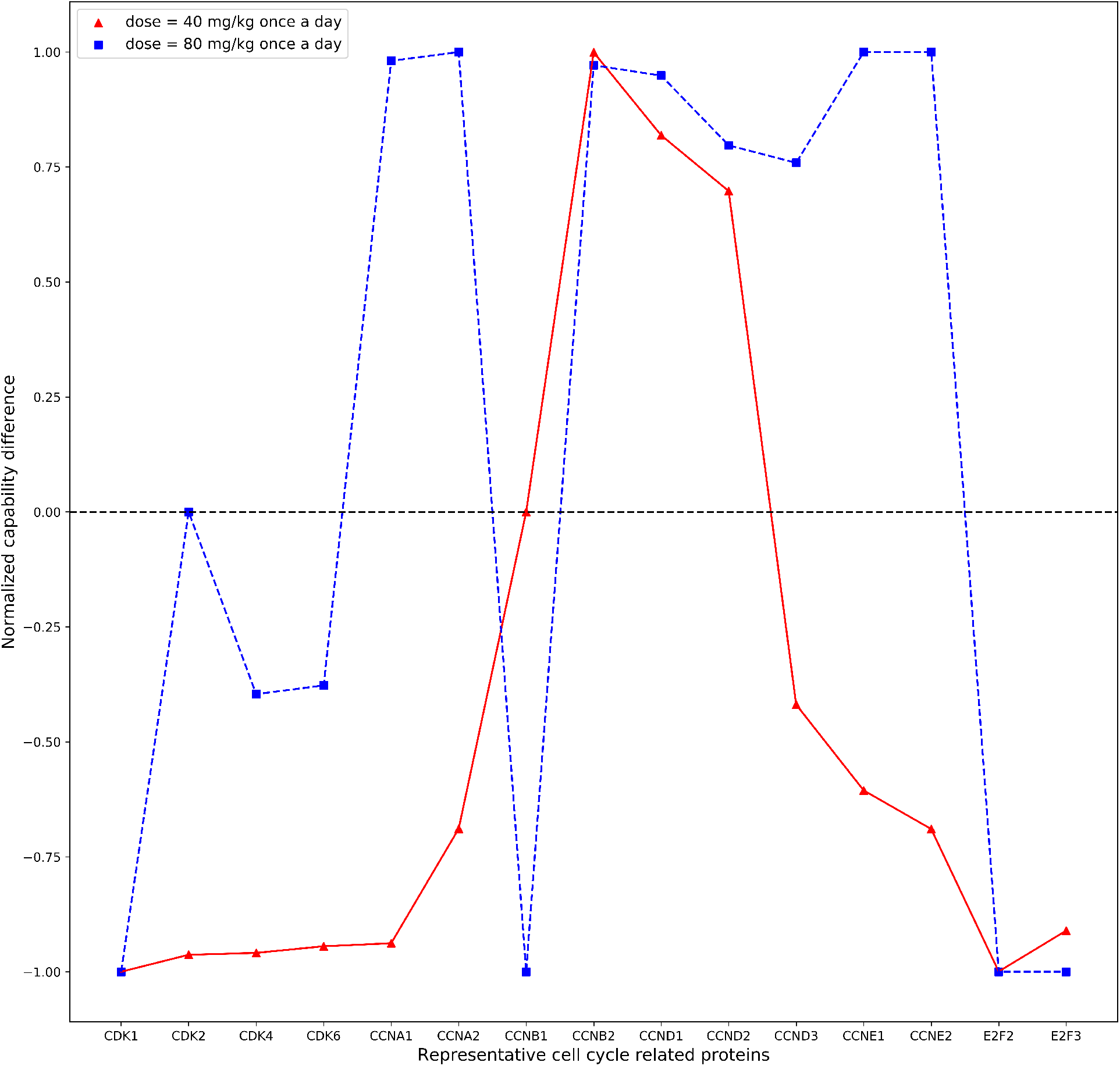
Normalized capability difference = [inhibition score − activation score] / max [inhibition score, activation score] between representative cell cycle related proteins.

