## Supplementary File for "A Signaling Network based Computational Model to Uncover Loop as the Novel Molecular Mechanisms for Medulloblastoma"

**Supplementary File S1: Literature validated therapeutic targets**

**Table 1**: literature confirmed therapeutic targets for WNT subtype MBs.

| **WNT type MBs** | **References** |
| --- | --- |
| PI3K/AKT | (1) |
| PP1 | (2) |
| PP2A |  |
| LRP5 | (3) |
| WNT3A |  |
| GSK3B | (3–5) |
| PARP | (6) |
| DKK1 | (7) |
| WIF1 |  |
| PPARG | (8) |

**Table 2**: literature confirmed therapeutic targets for SHH subtype MBs.

| **SHH Type MBs** | **References** |
| --- | --- |
| PI3K | (9) |
| MTOR |  |
| AKT |  |
| IKBKB |  |
| JUN |  |
| CCND1 |  |
| ERK |  |
| CDKN1A |  |
| MEK | (10) |
| PRKACA |  |
| YAP1 | (11) |
| IRS1 |  |
| EGFR | (12) |
| PRKCA |  |
| SMO |  |
| HDAC |  |

**Table 3**: literature confirmed therapeutic targets for Group 3 subtype MBs.

| **Group 3 type MBs** | **References** |
| --- | --- |
| CDK1 | (13) |
| GSK3B |  |
| CDK2 | (14) |
| STAT3 | (15) |
| TP53 | (16) |
| MYC |  |
| PI3K | (17) |
| PP2A |  |
| IGF |  |
| MTOR | (18) |
| PTEN | (19) |
| KDR | (20) |
| RAS | (21) |
| JNK |  |
| RAF |  |
| MEK |  |
| ERK |  |
| XIAP |  |
| RAC1 |  |
| CSNK1E |  |
| CDK4 |  |
| SRC |  |
| CAM Kinase |  |
| protein kinase A |  |
| protein kinase C |  |
| MAPKAPK2 |  |
| EGFR |  |
| PDPK1 |  |
| AKT |  |
| BCL2 |  |
| IKBKB |  |
| TNF |  |
| NFKB1 |  |
| SHH |  |
| PPP3R2 |  |
| HSP90 |  |
| IGF2 |  |
| WNT |  |
| ADORA3 |  |
| HDAC |  |
| CSNK1G1,2,3 |  |
| HTR1A |  |
| ARBB8 |  |
| PPARG |  |
| ADRA1A,B,D |  |
